# Exploring symbiotic legume-rhizobia relationships across tropical species

**DOI:** 10.1101/2024.08.23.609460

**Authors:** Keilor Rojas-Jimenez, Jéssica Morera-Huertas, Marianne de Bedout, Beatriz Rivera, Andrés Zúñiga, Jose A. Molina-Mora, Laura Solís-Ramos, Mario A. Blanco, Oscar J. Valverde-Barrantes

## Abstract

- The nitrogen-fixing symbiosis with rhizobia significantly contributes to the successful establishment of legumes in tropical environments. Here, we explored the symbiotic interactions between legumes and rhizobia across 109 tropical species.
- We evaluated the presence of root nodules in 72 genera spanning four Fabaceae subfamilies, encompassing 53% of the genera in Costa Rica and approximately 9.4% globally. Our analysis revealed root nodules in 78% of the species belonging to Caesalpinioideae (67%) and Papilionoideae (84%). Also, we formulated a predictive model for nodulation in legumes, with fine-root color and flower symmetry as the main predictor criteria.
- Nodulating taxa consistently presented higher N levels in tissues, thinner roots, and a pronounced phylogenetic influence on nodulation. We also identified the bacterial genera associated with all nodulating legumes. Caesalpinioideae formed symbiotic relationships with ten rhizobial genera, whereas Papilionoideae were associated with only four. *Bradyrhizobium* emerged as the predominant symbiont in nodules, while *Rhizobium*, *Mesorhizobium*, *Agrobacterium*, *Paraburkholderia*, and *Cupriavidus* exhibited a more restricted host range. Furthermore, genomic analysis suggests that 15 out of the 22 sequenced rhizobial strains may represent novel species.
- Our study enhances the understanding of the different dimensions that comprise legume-rhizobia interactions, incorporating a diverse array of previously unexamined tropical forest legume taxa.

## Introduction

While the mutually beneficial partnership between nitrogen-fixing bacteria (rhizobia) and legume plants is relatively well-studied in agricultural settings, there’s a significant gap in knowledge of natural forests. This lack of understanding exists even though this type of symbiosis appears to be crucial for the dominance of Fabaceae in tropical ecosystems (Burnham & Johnson, 2004; Parker, 2008; Gei *et al*., 2018; de Bedout-Mora *et al*., 2022; Mathesius, 2022).

Nodulation provides legumes with essential nitrogen, a plant macronutrient and key element for growth and reproduction, helping legumes thrive and diversify in ecosystems where nitrogen is scarce (Doyle, 2016; Sprent *et al*., 2017; James, 2022). The interaction manifests itself through the development of root nodules that intracellularly host the nitrogen-fixing bacteria (Griesmann *et al*., 2018; Ferguson *et al*., 2019a; de Faria *et al*., 2022). There, the bacteria convert atmospheric nitrogen, unusable by plants, into a usable form such as ammonium, and deliver it to the host. Therefore, nitrogen-fixing leguminous plants hold a substantial competitive advantage in acquiring this essential nutrient compared to non-fixing species This process not only benefits the legumes themselves but also enriches the ecosystem by incorporating nitrogen throughout the food chains (Sprent, 2008b; Ferguson *et al*., 2010; Adams *et al*., 2016; Ferguson *et al*., 2019b).

The rhizobial group comprises nitrogen-fixing Gram-negative bacteria that are aerobic and heterotrophic. They belong to the Alphaproteobacteria and Betaproteobacteria classes including genera such as *Rhizobium*, *Bradyrhizobium*, *Agrobacterium*, *Allorhizobium*, *Mesorhizobium*, *Ensifer*, *Cupriavidus*, and *Ralstonia* (Vessey *et al*., 2005; Sprent, 2008a; Doyle, 2016). Some rhizobial strains exhibit a broad host range, allowing them to form nodules and fix nitrogen in numerous species (Bala & Giller, 2001; Sprent, 2008b), while others rely on genes encoded in plasmids to establish symbiosis and fix nitrogen (Roumiantseva & Muntyan, 2015).

Legumes carefully manage the trade-off between acquiring nitrogen and allocating resources toward its fixation (Ferguson *et al*., 2010, 2019b). The process is initiated by the legume secreting flavonoids into the rhizosphere to attract rhizobia. The flavonoids induce the bacteria to release nod factors, which in turn trigger the transcriptomic reprogramming of the host to initiate nodulation, involving the coordinated function of more than 30 genes (Ferguson & Mathesius, 2014; Sprent *et al*., 2017; Griesmann *et al*., 2018; Roy *et al*., 2020; Yang *et al*., 2022). However, it is the host, who ultimately accepts or rejects the bacterial invitation to establish the symbiotic association (Tian *et al*., 2013; Ferguson *et al*., 2019b).

The ability to form nodules in the Fabaceae is not evenly distributed among its species. Based on the latest classification of the family (Azani *et al*. 2017), the formation of nodules is phylogenetically limited to many (but not all) members of the two largest subfamilies (Papilionoideae and Caesalpinioideae), whilst the four other subfamilies (Cercidoideae, Detarioideae, Dialioideae, and Duparquetioideae) lack known nodulating members (Werner *et al*., 2014; Li *et al*., 2015; Tedersoo *et al*., 2018). Albeit this lack of nodule formation seems to be associated with the presence of specialized active genes in the plants (Zhao *et al*., 2021), little information has been collected about morphological traits in roots that may condition their colonization by nitrogen-fixing bacteria. However, recent developments in the identification of strategies adopted by roots to acquire nutrients suggest that variations in root morphology and chemistry could be important factors determining the dependency of plants to maintain symbiotic associations belowground (Xia *et al*., 2021a; Weigelt *et al*., 2021).

Previous studies have explored the ability of certain tropical leguminous tree species to form root nodules, with some successfully identifying the rhizobial symbionts (Ramírez & Flores, 1994; Budowski & Russo, 1997; Menge & Chazdon, 2016; Gei *et al*., 2018; de Bedout-Mora *et al*., 2022). However, several important knowledge gaps persist regarding the possible co-evolution between rhizobial groups and legume clades, as well as determining the ability of nodulation in many legume taxa at lower taxonomic levels, the specific rhizobial species involved in symbiosis, and their host range. Additionally, there is interest in exploring the feasibility of establishing predictive models for nodulation based on morphological plant traits (including those of roots, leaves, and flowers).

Part of the limitation of the information relies on the relatively poor phylogenetic representation of the clades studied so far. Most studies focused on symbiotic interactions between Fabaceae and rhizobial bacteria are restricted to lab studies with model plants (often agricultural crops), ignoring thousands of wild species of minor or no economic importance. In other cases, it is indicated whether a species of legume can nodulate, but it is not detailed with whom it establishes the symbiotic relationship. This situation has created a gap of information regarding the symbiotic affiliations between host and microbial communities, limiting our ability to understand the evolutionary patterns between bacteria and plants at large.

Costa Rica stands out for its remarkable biological diversity, particularly in the realm of legumes. The country boasts at least 136 legume genera and 593 species, including representatives from five of the six subfamilies of Fabaceae. Furthermore, Fabaceae boasts more tree species than any other plant family in Costa Rica, with 178 out of the 1407 tree species (12.6%) belonging to this family (Zamora, 2010; Programa REDD/CCAD-GIZ-SINAC, 2015). Thus, Costa Rica offers an excellent opportunity for investigating symbiotic relationships between tropical legumes and their associated rhizobial symbionts. These data also highlight the strategic importance of studying the potential contribution and significance of symbiosis with nitrogen-fixing bacteria to the ecological success of legumes in tropical environments (Menge & Chazdon, 2016; Taylor *et al*., 2017; Gei *et al*., 2018).

In this study, we investigated 109 tropical legume species from four subfamilies of Fabaceae. Our analysis involved assessing nodulation presence, isolating and characterizing rhizobial endosymbionts, and examining the associations between nodulating capacity and plant morphological and anatomical traits. The insights derived from this research are particularly valuable as they help fill knowledge gaps regarding the relationships between legumes and rhizobia, especially in tropical ecosystems, which have received comparatively less attention.

## Materials and Methods

### Plant material and root nodule assessment

In this study, we investigated 109 tropical legume species from four subfamilies of Fabaceae. The trees used in this study were obtained from tree nurseries in different locations of Costa Rica (Vivero Forestal EDECA, Universidad Nacional; CODEFORSA, San Carlos; Centro de Conservation Santa Ana; Osa Conservation; Corredor Biológico Talamanca Caribe) representing species from different ecosystems in the country, including the Caribbean rainforest, Pacific seasonal dry forest, and Central Valley mid-elevation ecosystems. The 109 species were grouped into 72 genera of four Fabaceae subfamilies (Caesalpinioideae, Cercidoideae, Detarioideae, and Papilionoideae), representing ca. 53% of the legume genera reported for Costa Rica and ca. 9.4% of the world legume genera (**Table 1**).

**Table 1.**
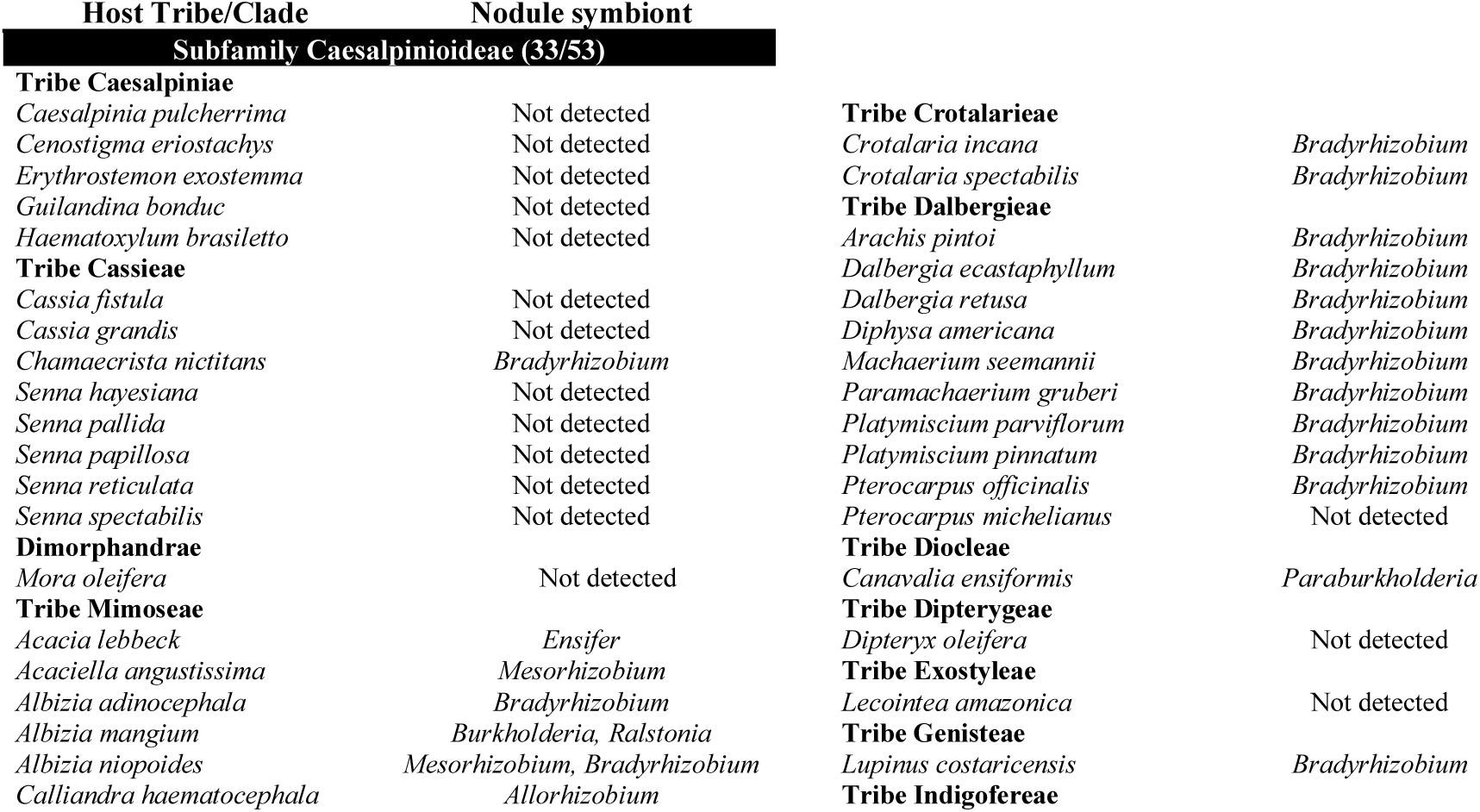
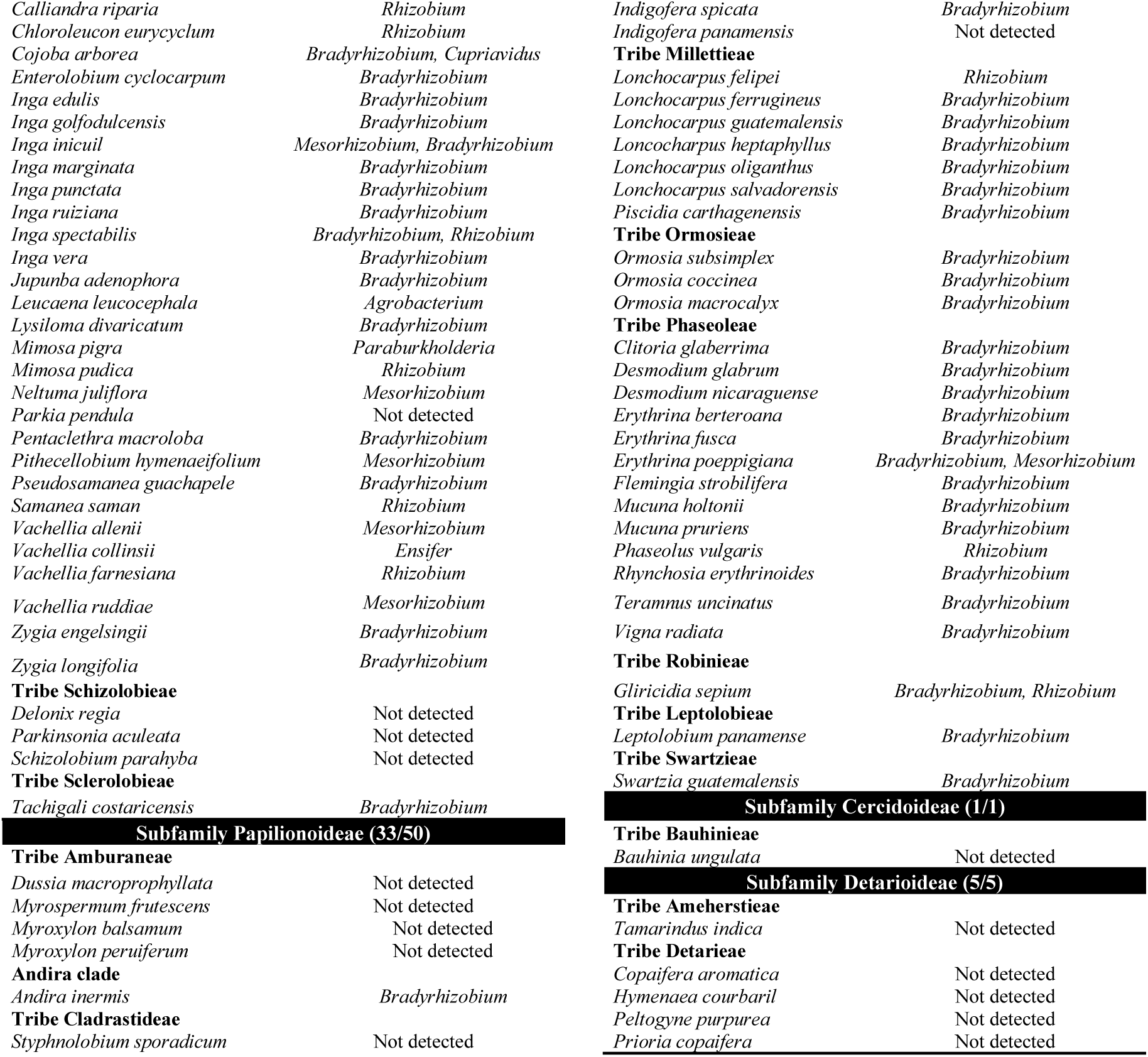
Nodulation status of the 109 tropical legume species examined in this study, categorized by their tribe/clade and Fabaceae subfamily (subfamilial classification follows The Legume Phylogeny Working Group 2017, by Azani et al. 2017; the tribal classification follows Bruneau et al. 2024 for Caesalpinioideae, Cardoso et al. 2013 for Papilionoideae and de la Estrella et al. 2018 for Detarioideae). The number of sampled genera and species within each subfamily is included in each subfamily heading. The asterisk (*) highlights threatened species. For all legume species exhibiting root nodules, we provide identification of their rhizobial symbionts to genus level (from both cultured strains and metabarcoding; see Materials and Methods), while species lacking nodules are indicated as “Not detected”. We employed both culture-dependent and culture-independent approaches to gather and compile this data.

For this work, we used small trees with 30-100 cm height and aged 1-3 years. The plants were acclimatized for 3-6 months in a common house garden located in Barva de Heredia, Costa Rica (1100 m elevation, 18-28°C diurnal temperature range). The plants were transferred from plastic bag containers to pots, using the same substrate where the plants were originally obtained. During the acclimatization period, the plants received daily irrigation without any fertilizer or pesticide application. Access permission VI-4056-2020 was obtained from the Biodiversity Commission of the University of Costa Rica.

To determine the presence of nodules, the roots of each plant were manually examined by removing the adhered soil. A qualitative determination of the coloration (light or dark) was recorded, the density of the roots (roots per square centimeter of soil) was estimated, and the size and shape of the nodules, if any were also recorded. A small segment of the root was pruned and washed with water. If the root presented nodules, a subsample was immediately taken to the laboratory for bacterial isolation (1-4 nodules depending on their size) and another subsample was preserved in mineral oil for subsequent analysis of root traits. Once sampling was done, each individual tree was replanted in a pot, kept in a nursery for approximately one year, and then planted on the ground for *ex-situ* conservation.

### Plant nodulation predictive model

(Breiman *et al*., 2017)In order to create a simple model that predicts whether a Fabaceae species forms nodules, we compiled a database with nodule presence as the dependent variable, and various plant characteristics as independent variables, including leaf type, fiotaxia, presence of terminal leaflet, leaflet position, flower color and flower symmetry, thorns presence, fine-root color, and taxonomic subfamily. We used machine learning, specifically the decision tree algorithm (Breiman *et al*., 2017), to create the model. The decision tree was generated in R using packages caret (Kuhn, 2008), rpart (Therneau & Atkinson, 2023), and rpart.plot (Milborrow, 2024).

### Bacterial isolation and DNA extraction from root nodules

Upon arrival in the laboratory, the nodules were transferred to 2 mL tubes and rinsed twice with sterile distilled water. Next, they underwent surface disinfection with 70% ethanol for 1 minute, followed by two additional rinses with sterile distilled water. To ensure sterility of external surfaces, the nodules were then treated with 2% sodium hypochlorite for 5 minutes, subsequently undergoing seven rinses with sterile distilled water. Following disinfection, the nodules were aseptically transferred to new tubes containing 250 μL of sterile Phosphate Buffered Saline (PBS, pH 7.4). The root nodules were then macerated using autoclaved plastic pistils to release the bacterial contents. The resulting suspension was serially diluted in tenfold increments (10^-1^ and 10^-2^) using PY broth (5 g/L peptone, 1 g/L anhydrous dextrose, 0.5 g/L dipotassium phosphate, 0.2 g/L magnesium sulfate, 0.1 g/L sodium chloride). Subsequently, 50 μL aliquots from each dilution were spread onto Petri dishes containing PY-Agar supplemented with cyclohexamide (40 mg/L) to suppress fungal growth. These plates were incubated for up to three weeks at 28°C until bacterial colonies were visually detectable. Individual colonies were then carefully picked and streaked onto fresh PY-Agar plates containing the same antifungal agents indicated above. Finally, the purified strains were cryopreserved in PY broth supplemented with 20% glycerol and stored at −80°C in the collection of the Laboratory of Genetics and Ecology of Microorganisms (LEGMi) at the School of Biology, University of Costa Rica.

### Extraction of DNA and molecular identification of plants and bacterial isolates

The identification of all legume species used was verified by botanical experts specializing in legume taxonomy. Additionally, molecular identification was performed using the *mat*K gene marker. For this step, approximately 250 mg of leaf tissue was collected in a 2 mL tube. The tissue was then frozen with liquid nitrogen and physically disrupted using a sterilized plastic pestle. DNA was extracted from the resulting material using the DNEasy PowerSoil Kit (Qiagen, USA) following the manufacturer’s instructions. Similarly, DNA extraction for bacterial isolates involved collecting a loopful of culture strain and resuspending it in 250 µL of PBS. The DNA was then extracted from this suspension using the same kit.

For plants, the plastid *mat*K gene was amplified using the primers matK472F and matK1248R (Yu *et al*., 2011). For bacteria, the 16S rRNA gene was amplified using primers 27F and 1492R (Lane, 1991). Amplification involved an initial denaturation at 98°C for 2 minutes, followed by 35 cycles of denaturation at 96°C for 30 seconds, annealing at 50°C (*mat*K) and 52°C (16S rRNA) for 30 seconds, and extension at 72°C for 40 seconds and a final extension at 72°C for 10 minutes. Amplification products were visualized on a 1% agarose gel. The amplicons were sequenced with Sanger technology at Macrogen (South Korea). Sequences were assembled and edited using BioEdit (Hall, 1999). Plant identifications were obtained by comparing their sequences with the NCBI’s nr database. Currently accepted scientific names were verified at the Plants of the World Online webserver (https://powo.science.kew.org/). Bacterial taxonomic classification was carried out by comparing sequences with reference databases SILVA v138.1, GTDB v214.1, and NCBI’s 16S rRNA database. Sequence data were deposited at the GenBank under accession numbers PP449130-PP449202 (for *mat*K) and PP439182-PP439285 (for the 16S rRNA).

For the phylogenetic analysis of plants and bacterial isolates belonging to families containing nitrogen-fixing symbionts such as Rhizobiaceae, Xanthobacteraceae, and Burkholderiacea, we aligned the respective sequence datasets with MAFFT v7 (Katoh *et al*., 2019). For nearly 20% of plant species, it was not possible to obtain *mat*K sequences, therefore corresponding sequences were obtained from Genbank. We generated a maximum likelihood phylogenetic tree with IQ-TREE web server (Trifinopoulos *et al*., 2016), using the best-fit model of evolution (TVM+F+G4 for *mat*K and TN+F+R3 for 16S rRNA). The resulting phylogenetic trees were displayed and edited in MEGA-X (Kumar *et al*., 2018).

### Metabarcoding of root nodules

Since culturing bacteria from nodules was not successful for some plant species, we also used DNA metabarcoding to identify the symbionts directly from the nodules. We also used this technique because we hypothesize that in each nodule it is possible to identify multiple species of diazotrophic bacteria. We extracted DNA using the DNEasy PowerSoil Kit (Qiagen, USA) from the superficially disinfected nodules that were macerated in PBS, as described previously. The prokaryotic amplicon library was constructed based on the V4 hypervariable region of the 16S rRNA gene using universal primers 515F and 806R (Caporaso *et al*., 2011). The PCR conditions consisted of 95°C for 3 min initial denaturation followed by 35 cycles at 95°C for 45 s, 53°C for 1 min, 72°C for 1 min, and a final extension at 72°C for 5 min. The PCR products with the proper size were selected by 1% agarose gel electrophoresis. Each sample’s exact amount of PCR products was pooled, end-repaired, A-tailed, and further ligated with Illumina adapters. Libraries were sequenced on a paired-end Illumina platform to generate 250 bp paired-end raw reads (Illumina Novaseq, Novogene Bioinformatics Technology Co., Ltd, CA, USA).

We used the DADA2 version 1.21 to process the Illumina-sequenced paired-end fastq files and to generate a table of amplicon sequence variants (ASVs), which are higher-resolution analogs of the traditional OTUs (Callahan *et al*., 2016). Briefly, we removed primers and adapters, inspected the quality profiles of the reads, filtered and trimmed sequences, estimated error rates, modeled, and corrected amplicon errors, and inferred the sequence variants. Then, we merged the forward and reverse reads, removed chimeras, constructed the ASV table, and assigned taxonomy. For the taxonomic assignment, we used the SILVA v138.1 reference database (Quast *et al*., 2013). We verified the consistency between the taxonomic assignments of the SILVA database and those of NCBI using the BLAST tool (Altschul *et al*., 1990). Sequence data were deposited at the NCBI Sequence Read Archive under accession number PRJNA1084876 (https://www.ncbi.nlm.nih.gov/sra/PRJNA1084876). In the dataset, we removed sequences assigned to Chloroplast and Eukarya. The average number of sequences per sample was 78033 (ranging from 17843 to 146702).

The statistical analyses and visualization of results were performed with the R statistical program (R Core Team, 2023) and the Rstudio interface. Package Vegan v2.6-4 (Oksanen *et al*., 2022) was used to calculate alpha diversity estimators and non-metric multidimensional scaling analyses (NMDS). Data tables with the amplicon sequence variants (ASV) abundances were normalized into relative abundances and converted into a Bray–Curtis similarity matrix. To determine significant differences between the bacterial community composition according to the legume subfamily, we used the non-parametric multivariate analysis of variance (PERMANOVA). To estimate differences in the relative proportion of microorganisms and the diversity indices, we used the non-parametric Kruskal-Wallis test, while pairwise comparisons were performed using Wilcox and Dunn’s tests (with Bonferroni adjustment).

### Whole genome sequencing and assembly

To improve the precision of the 16S rRNA taxonomic assignments and explore potential novel rhizobial species, we selected 22 strains representing diverse branches of the phylogenetic tree previously generated. These strains underwent whole-genome sequencing after re-culturing, and DNA extraction with the DNEasy PowerSoil Kit (Qiagen, USA).

Sequencing was performed at Novogene Co. (CA, USA) using a NovaSeq PE150 sequencing platform (Illumina, Inc.). Genome assembly analyses were performed based on the bioinformatic pipeline by (Molina-Mora *et al*., 2020). The raw sequencing data underwent quality assessment and filtering steps using FastQC (Andrews, 2010) and Trimmomatic (Quality score >30) (Bolger *et al*., 2014), respectively. Taxonomic identity analysis was conducted using average nucleotide identity (Ciufo *et al*., 2018) through the Read Assembly and Annotation Pipeline Tool (https://www.ncbi.nlm.nih.gov/rapt/). Genome assembly was accomplished using the Unicycler algorithm (Wick *et al*., 2017), and its quality was evaluated based on the 3C criterion (contiguity, completeness, and correctness) (Molina-Mora *et al*., 2020), incorporating results from QUAST (Gurevich *et al*., 2013) and the BUSCO tool (Simão *et al*., 2015).

### Root traits and chemical analyses

To identify root morphological characteristics associated with the nodulation phenotype in tropical legumes, we analyzed root traits across a subset of 25 species, comprising 15 nodulating and 10 non-nodulating ones. We collected intact and well-branched woody fine-root segments, each measuring 5-10 cm in length comprising the first three root orders.

These segments were thoroughly washed and preserved in mineral oil until processing. In the laboratory, the mineral oil was removed, and the roots were laid out in a transparent tray submerged in water. Subsequently, they were scanned at 600 dpi resolution to quantify various parameters, including root average diameter, total length, branch points, branch frequency, root tips, and total length, utilizing RhizoVision Explorer (Seethepalli *et al*., 2021). Furthermore, the nitrogen content, expressed as a percentage of dry weight, was assessed in a subset of 12 nodulating and non-nodulating legume species, encompassing separately leaf and root tissues. The determination of nitrogen content was conducted using atomic absorption spectroscopy.

## Results

Of the 109 legume species examined in this study, 53 belong to subfamily Caesalpinioideae, 50 to subfamily Papilionoideae, five to subfamily Detarioideae, and one to subfamily Cercidoideae (**Fig. 1**, **Table 1**). Our analysis revealed root nodules in 78% of the species. However, a notable disparity exists among subfamilies. No nodulation was observed in the Cercidoideae and Detarioideae subfamilies. In contrast, 67% of Caesalpinioideae species exhibited nodules, while Papilionoideae displayed the highest proportion at 84%.

**Fig. 1.**
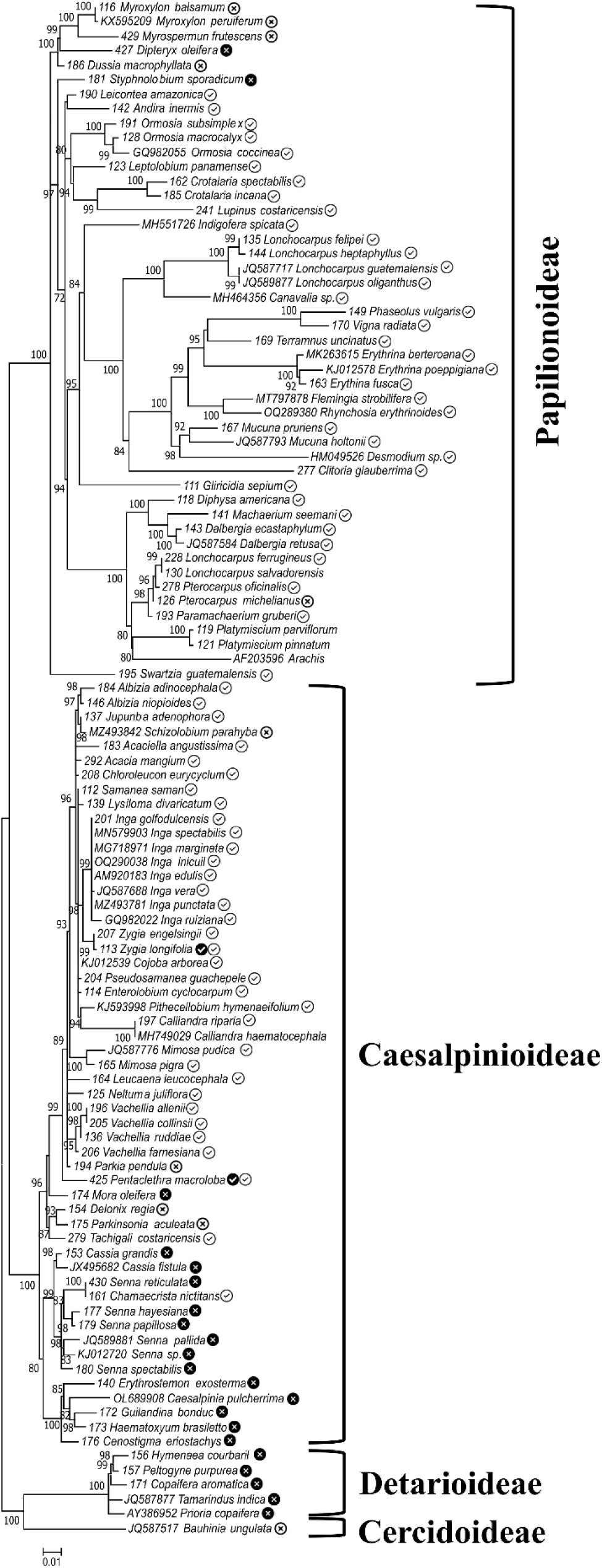
Phylogenetic relationships of the legume species examined in this study, based on *mat*K sequence data. Sequences were aligned using MAFFT, and maximum likelihood phylogenetic trees were constructed using IQ-TREE, employing the best-fit models of evolution (TVM+F+G4). The resulting tree was visualized and edited in MEGA-X, with bootstrap values depicted on the nodes. Dark or light circles next to each species show whether the fine roots are light or dark colored, respectively; symbols in circles show whether the species formed nodules (✓) or not (**×**).

By using machine learning tools, we developed a model (Accuracy = 96%) that can be used to predict whether a Fabaceae species will nodulate, relying solely on two variables: fine-root color and flower symmetry. It works as follows: if the species displays light-colored roots, there is a very high probability it will form nodules. If it showcases dark roots alongside actinomorphic flowers, the probability of nodulation is high. However, if it features dark roots and zygomorphic flowers, the likelihood of forming nodules is null (**Fig. 2**).

**Fig. 2.**
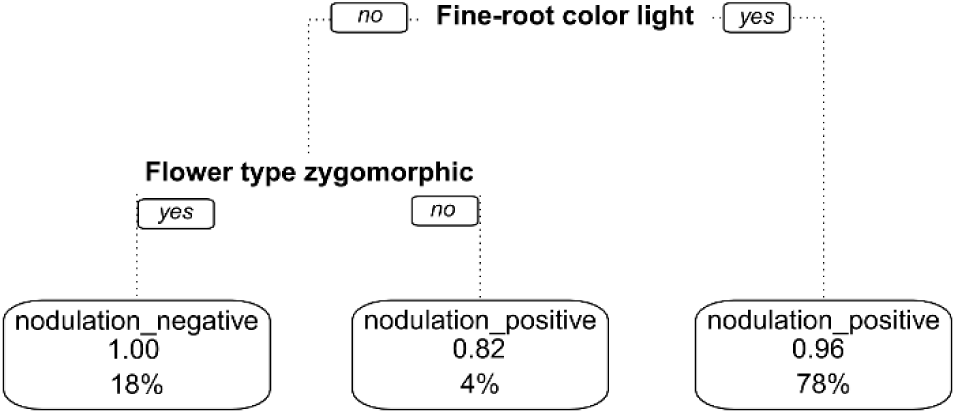
Decision tree model to predict whether a Fabaceae species forms root nodules based on two characteristics: fine-root color and flower symmetry. According to this model (Accuracy = 96%), if the fine roots are light-colored, the species has a probability of 0.96 of presenting nodules. If the roots are dark and the flowers are actinomorphic, the probability is 0.82, but if the roots are dark and the flowers are zygomorphic, the probability of not forming nodules is 1.00. The percentages represent the amount of data (proportion of sampled species) in each category.

Among the plants that form nodules, we were able to isolate a large diversity of rhizobia, based on the analysis of the 16S rRNA gene sequences. The 104 rhizobial strains isolated belonged to four taxonomic families and seven genera (**Fig. 3**). The most abundant genera were *Bradyrhizobium* and *Rhizobium*. Notably, we also isolated and identified the genera *Burkholderia*, *Paraburkholderia*, and *Cupriavidus* of Betaproteobacteria. More detailed information on the phylogenetic relationships of the bacterial isolates obtained is shown in **Supplementary Fig. 1**.

**Fig. 3.**
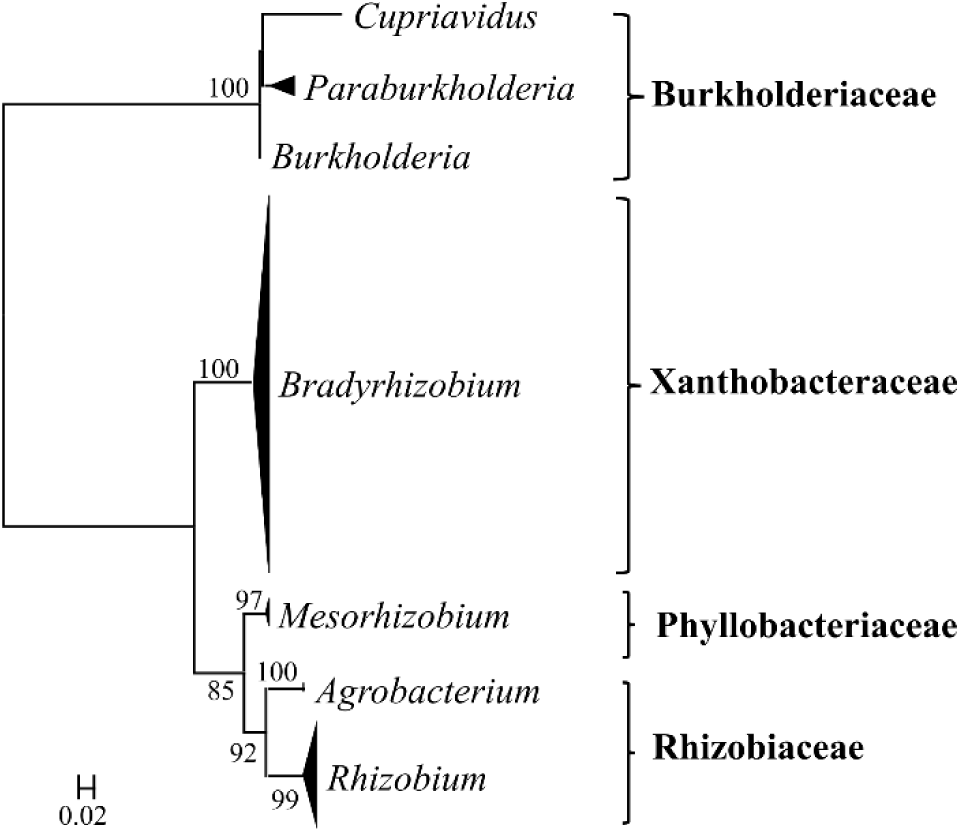
Phylogenetic relationships of the cultured rhizobial strains isolated in this study, based on 16S rRNA sequence data, grouped by taxonomic genus and family. The 16S rRNA sequence dataset was aligned with MAFFT, and the maximum likelihood phylogenetic tree was constructed using IQ-TREE, employing the best-fit models of evolution (TN+F+R3). The resulting trees were visualized and edited in MEGA-X, with bootstrap values depicted on the nodes.

Since it was not possible to obtain isolates from some legume species, we also examined the nodule rhizobial microbiome using DNA from the superficially disinfected root nodules. Specifically, we analyzed the nodule microbiome of 40 legume species belonging to Caesalpinioideae (n = 16) and Papilionoideae (n = 24). This approach enabled us to assess the composition of bacterial communities within the nodules and discern any notable distinctions between the two subfamilies (**Fig. 4**).

**Fig. 4.**
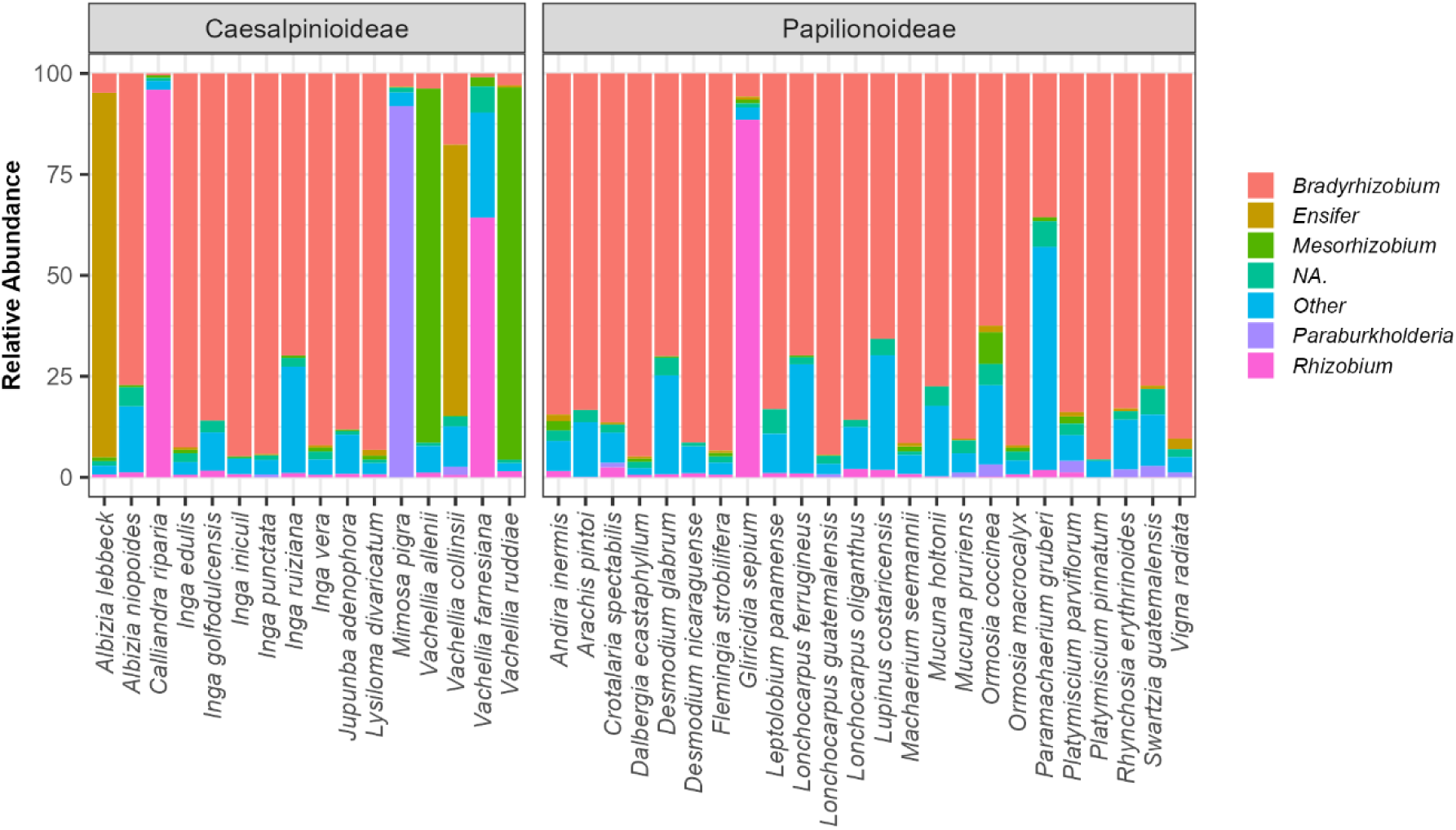
Rhizobial community composition, based on metabarcoding of the 16S rRNA gene, in root nodules of 40 tropical legume species belonging to the subfamilies Caesalpinioideae and Papilionoideae.

The prokaryote community within the nodules was composed of 6443 ASVs, according to sequence analysis of the V4 region of the 16S rRNA gene. These sequences were assigned to 53 phyla and 127 classes. Proteobacteria was the most prevalent group, representing 92% of all sequences. Despite the presence of diverse species, nitrogen-fixing bacteria, particularly those belonging to the genus *Bradyrhizobium*, were most abundant across both Caesalpinioideae and Papilionoideae subfamilies. Members of *Rhizobium*, *Ensifer*, *Mesorhizobium,* and *Paraburkholderia* were also identified but within a more restricted host range. It was even possible to observe different rhizobial species coexisting inside the nodules. Also, a noteworthy case is that of genus *Vachellia*, where nodules from different species hosted distinct genera of nitrogen-fixing bacteria including *Mesorhizobium*, *Ensifer* and *Rhizobium*.

Alpha diversity analyses of the microbial communities in the nodules of Caesalpinioideae and Papilionoideae determined high values for species richness in these structures. On average, 295 ASVs per individual were found, ranging from 96 to 897 ASVs (**Supplementary Fig. 2**). Likewise, the average value of the Shannon Index was 1.5, with a range from 0.5 to 3.13. However, statistical analysis did not find significant differences in the values of diversity indices between both subfamilies studied.

The NMDS analysis revealed a strong dominance of nitrogen-fixing bacteria on the bacterial community structure within the nodules. This resulted in distinguishable clusters. Statistical tests further confirmed significant variations (Permanova, p=0.001) in the microbial community composition between Caesalpinioideae and Papilionoideae nodules (**Fig. 5**). Notably, Papilionoideae nodules harbored only two dominant nitrogen-fixing genera, whereas Caesalpinioideae nodules harbored five genera of these bacteria.

**Fig. 5.**
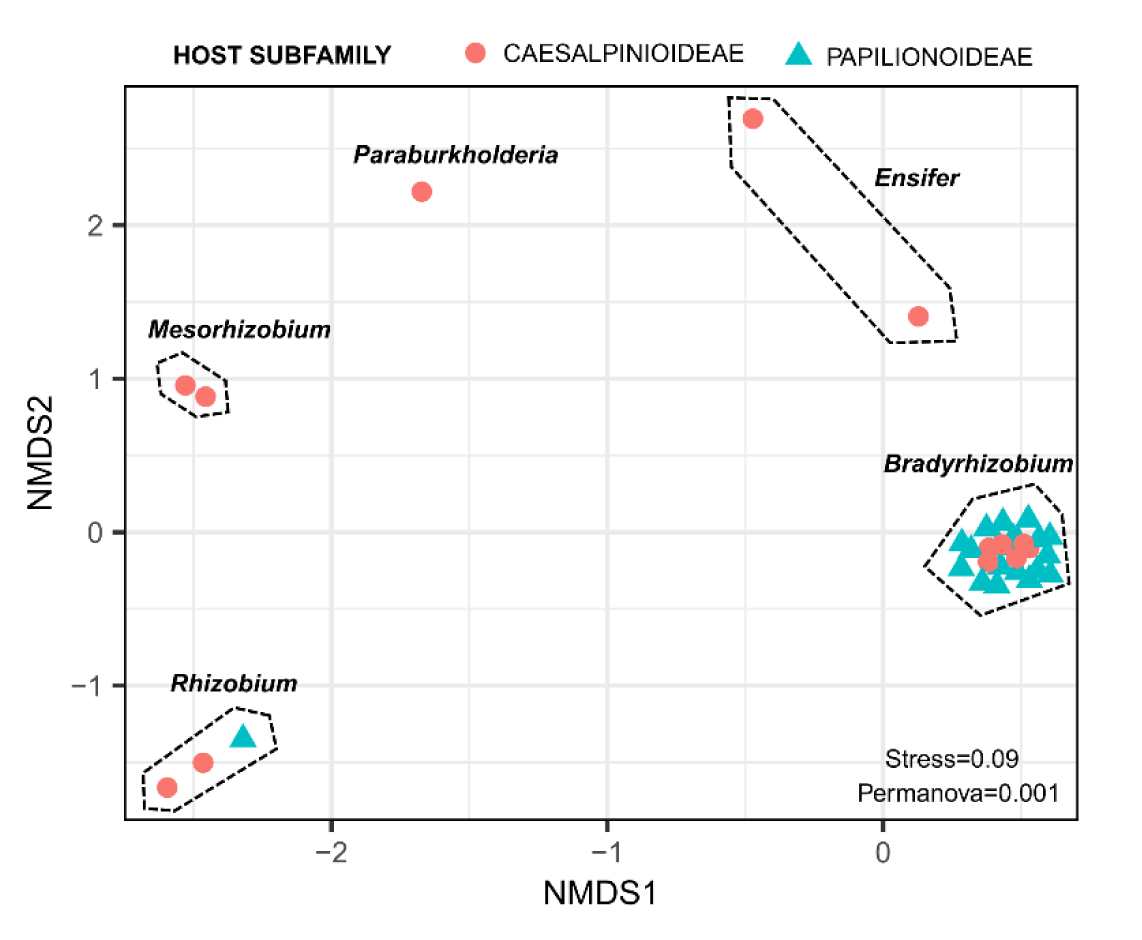
NMDS analysis of microbial community structure in root nodules of tropical legumes belonging to the subfamilies Caesalpinioideae and Papilionoideae. Clusters are marked with dashed lines and the dominant nitrogen-fixing bacterial genus in each cluster is shown.

To better understand certain characteristics associated with nodulation phenotype, we evaluated functional root traits of 25 species belonging to three subfamilies of Fabaceae (**Fig. 6A**). Among the variables measured, it was notable that the average root diameter in nodulating species was significantly smaller compared to non-nodulating species (Wilcoxon-Mann-Whitney test, p=0.003).

**Fig. 6.**
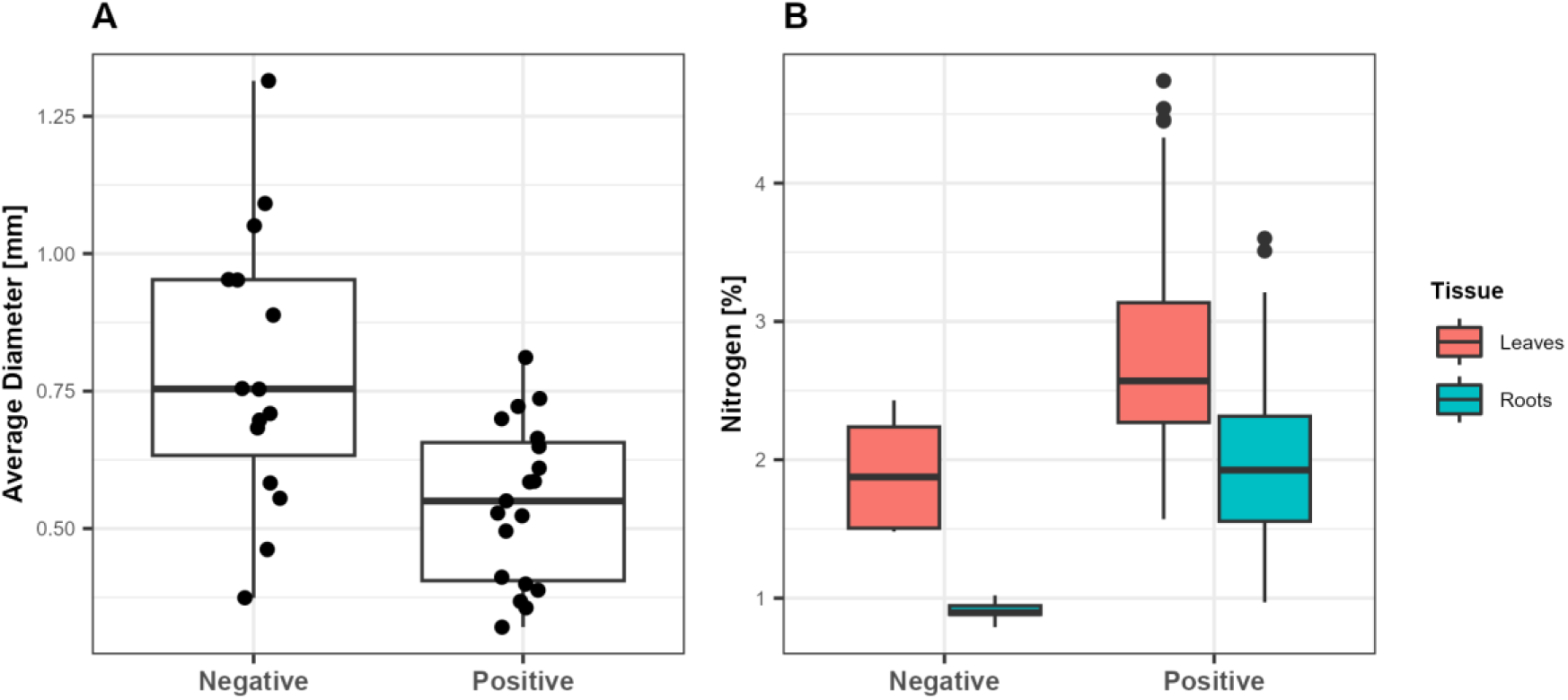
Average root diameter (**A**) and nitrogen content in both leaves and roots (**B**) of tropical leguminous plants are categorized by their nodulation capacity. Root diameter measurements were conducted across a sample of 25 species. The average root diameter in nodulating species was significantly smaller compared to non-nodulating species (Wilcoxon-Mann-Whitney test, p=0.003). The nitrogen content, assessed in a subset of 12 species, was significantly higher leaves and roots of nodulating species (Wilcoxon-Mann-Whitney, p<0.001) than in non-nodulating species.

Interestingly, variations between subfamilies were not evident. In addition, chemical analyses showed that the nitrogen content in both leaves and roots of nodulating species was significantly higher (Wilcoxon-Mann-Whitney, p<0.001) than in non-nodulating species (**Fig. 6B**). Specifically, we detected a higher nitrogen content in leaves (2.8% vs. 1.9%) and roots (2.0% vs. 0.9%) of nodulating plants compared to their non-nodulating species.

By integrating both culture-dependent and culture-independent techniques, we were able to identify the genera of the rhizobial symbionts associated with all nodulating legumes examined (**Table 1**). This analysis revealed a prominent phylogenetic influence on nodulation ability, particularly at the tribe level. Essentially, within the same legume tribe, either all species formed nodules or none did (for example, all of the sampled members of tribes Millettieae, Ormosieae, and Phaseoleae produced nodules; but none of the sampled members of tribes Amburaneae, Caesalpinieae, and Schizolobieae did). Only a few exceptions noted; for instance, within tribe Cassieae, *Chamaecrista nictitans* exhibited nodulation but species of *Cassia* and *Senna* did not, while *Parkia* did not exhibit nodules, but all other genera within tribe Mimoseae produced nodules.

Among the 36 species of Caesalpinioideae with root nodules, we detected ten rhizobial genera. In contrast, the 41 species of Papilionoideae with nodules were associated with only four rhizobial genera. Another noteworthy finding was the prevalence of the genus *Bradyrhizobium* as the dominant symbiont in both subfamilies. Other symbiotic genera like *Rhizobium*, *Mesorhizobium*, *Agrobacterium*, *Paraburkholderia*, and *Cupriavidus* displayed a more limited host range. Certain genera such as *Ensifer* were exclusively detected through metabarcoding. Interestingly, some host species were found to be colonized by different genera of rhizobia, typically involving *Bradyrhizobium* paired with another genus.

Finally, we conducted genome sequencing on 22 rhizobial strains belonging to different branches of the above-generated 16S rRNA phylogenetic tree. The genomic analysis confirmed the high quality of the resulting assemblies, considering metrics such as completeness, NS50, L50, and the number of contigs (**Table 2**). The most relevant result was that 15 out of the 22 strains displayed average nucleotide identity (ANI) values below 96% compared to the closest reference species, which is the threshold cut-off for defining species boundaries (Ciufo *et al*., 2018; de Lajudie *et al*., 2019). Consequently, these 15 strains represent candidates for the description of novel species. The formal characterization of these newfound species will be detailed in forthcoming publications.

**Table 2.**
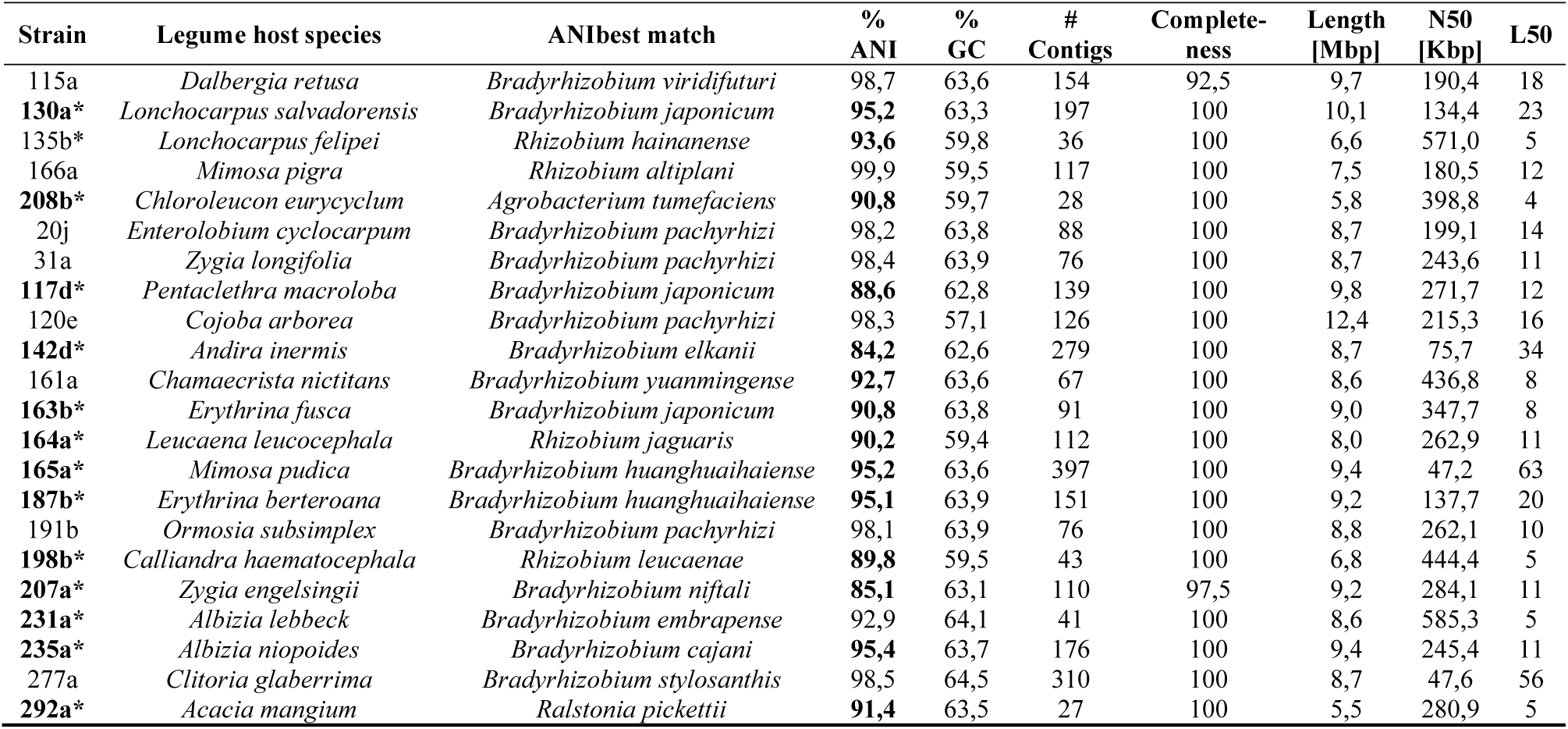
Quality analysis of the assembly and taxonomic identification of whole-genome sequences derived from 22 rhizobial strains isolated from tropical legumes. Strains with average nucleotide identity (ANI) values below 96%, indicating potential new species, are highlighted with an asterisk and in bold.

## Discussion

In our study, we unveil significant insights into the dynamics of legume-rhizobia relationships, particularly within tropical forest plants. We introduce a user-friendly method for predicting legume nodulation based on easily observable traits like root color and flower symmetry. We also delineate the contrasting nodulation proportions between Caesalpinioideae and Papilionoideae. Most importantly, we successfully identified the specific rhizobial bacteria associated with each of the legume species analyzed, providing important insights into these symbiotic partnerships.

Our findings align with prior research, reaffirming the absence of nodulation in the subfamilies Cercidoideae and Detarioideae, varying levels in Caesalpinioideae depending on the clade, and higher occurrences in Papilionoideae, excluding the ADA clade (Allen & Allen, 1981; Werner *et al*., 2014; Li *et al*., 2015; Tedersoo *et al*., 2018; de Faria *et al*., 2022). While most studies have primarily focused on recording presence or absence (and sometimes type) of nodulation, our research extends further by identifying the bacterial genera involved in the symbiotic relationship. We offer taxonomic details on the symbiont bacteria for approximately 12% of nodulating Fabaceae genera globally, according to the NodDB database (Tedersoo *et al*., 2018). A modified version of the NodDB database with symbionts is provided as **Supplementary File 1.**

The pronounced phylogenetic influence of the sampled legume species on nodulation ability is consistent with previous research, suggesting that species within the same tribe tend to uniformly either form root nodules or lack this capacity altogether. Our study reaffirms this trend and also agrees with the few exceptions within those groups (Werner *et al*., 2014; van Velzen *et al*., 2019; Zhao *et al*., 2021; de Faria *et al*., 2022). In addition, the absence of nodulation found within Cercidoideae and Detarioideae, and the presence of nodules in the Caesalpinioideae and Papilionoideae could be consistent with the “single gain and massive loss” hypothesis (Soltis *et al*., 1995; Werner *et al*., 2014; Griesmann *et al*., 2018; van Velzen *et al*., 2019; Geurts & Huisman, 2024).

Regarding the specific exceptions to uniformity within clades, our findings also align with previous observations indicating the lack of nodulation among species of tribe Cassieae, except for *Chamaecrista*, and the widespread capacity for nodulation in the large tribe Mimoseae (formerly treated as subfamily Mimosoideae), except in *Parkia*. Our results also show that the majority of Papilionoideae species are capable of forming nodules, except for members of the basal ADA clade (tribes Angylocalyceae, Dipterygeae and Amburaneae, with species of *Dipteryx*, *Myrospermum* and *Myroxylon* sampled by us, belonging to the last two tribes), as well as the Clasdrastideae and Exostyleae tribes (Allen & Allen, 1981; van Velzen *et al*., 2019; Zhao *et al*., 2021; Rahimlou *et al*., 2021). The nodulation ability observed in *Chamaecrista* has even been proposed as indicative of the genus’s pivotal role in the evolution of nodulation within Caesalpinioideae (Naisbitt *et al*., 1992; Sprent *et al*., 2013; de Faria *et al*., 2022; Braz *et al*., 2024).

Previous studies indicate that Fabaceae emerged around 4 million years before the Cretaceous-Paleogene mass extinction event roughly 65 million years ago. Following this event, the family experienced a rapid diversification, marked by numerous instances of whole genome duplication or triplication (Koenen *et al*., 2021; Zhao *et al*., 2021). This diversification primarily occurred within subfamilies Caesalpinioideae and Papilionoideae, and involved the duplication of specific genes and their refunctionalization; e.g., by N-terminal deletions in genes related to symbiosis with rhizobia such as NIN, RGP, NFR1, which play roles in recognizing Nod factors and facilitating the development of the infection thread during nodule formation (Griesmann *et al*., 2018; van Velzen *et al*., 2019; Mathesius, 2022; Geurts & Huisman, 2024).

According to this hypothesis, some losses or modifications in those genes subsequently occurred exemplified by changes observed in the Cassieae tribe of Caesalpinioideae and the ADA and Cladrastideae clades in Papilionoideae (van Velzen *et al*., 2019; Zhao *et al*., 2021). Subsequently, the latter subfamily underwent what is referred to as the 50 kb inversion, along with additional whole genome duplication processes, leading to even more significant diversification and the widespread maintenance of symbiosis with rhizobia. These factors may contribute to why Papilionoideae stands out as the most diverse group within Fabaceae and also exhibits the highest percentage of nodulation, as reaffirmed in this study.

However, we think that the loss of nodulation capacity should be considered beyond genetics and take into account ecological and evolutionary factors. For instance, the roots of tribe Cassieae, a highly prevalent group in tropical regions but mostly lacking in nodulation, exhibit a distinct phenotype characterized by dark (sometimes also green-colored), thin, and dense roots. This pattern contrasts with the majority of nodule-forming legumes, which typically display lighter-colored roots. Interestingly, this phenotypic disparity suggests a potential adaptive advantage, possibly sacrificing symbiosis to enhance defense mechanisms against pathogens. This adaptation could involve the production of substances like phenols, tannins, and other compounds that confer dark coloration (Sprent & James, 2007; Cheynier *et al*., 2013; Downie, 2014; Xia *et al*., 2021b).

Alternatively, there might be an energetic trade-off at play. That is, from an energetic perspective, it may be more efficient to invest resources in producing new roots annually rather than sustaining nitrogen fixation, a process that demands significant energy expenditure (Hoffman *et al*., 2014; Postma *et al*., 2014; Mathesius, 2022). In this regard, it is necessary to deepen on how the anatomical and even architectural features of the roots could be related to the establishment of symbiotic relationships with nitrogen-fixing bacteria.

Conversely, non-nodulating species within the ADA clade of Papilionoideae, as well as Detarioideae, exhibited thicker roots with a lower branching frequency when compared to nodulating groups. This observation might indicate a trade-off favoring the maintenance of durable thicker roots over time, contrasting with thinner roots that are possibly more seasonal (Gaitán-Alvarez *et al*., 2020). Alternatively, this root anatomy could facilitate symbiotic relationships with fungi and bacteria, potentially aiding in fulfilling nutritional requirements. However, these hypotheses require further investigation.

We initially expected an association between certain Fabaceae subfamilies and particular rhizobia genera, for instance, that *Bradyrhizobium* might predominate in Caesalpinioideae while *Rhizobium* in Papilionoideae (Martínez-Romero, 2009). However, our results contradict this hypothesis since *Bradyrhizobium* emerged as the dominant genus in most nodulating legumes, irrespective of the subfamily, a finding consistent across both culture-dependent and culture-independent techniques. Nonetheless, a closer examination at the species level suggests some associations, for instance, *B. cajani* exclusively occurred in plants of the Papilionoideae subfamily, whereas *B. ganzhouense* was confined to Caesalpinioideae.

It’s noteworthy that we identified ten rhizobia genera forming nodules in Caesalpinioideae, compared to four in Papilionoideae. Additionally, only genera from Betaproteobacteria were isolated from Caesalpinioideae. These findings support the idea that initial symbiotic events likely took place in this legume subfamily, potentially involving Betaproteobacteria (Brea *et al*., 2008; Bontemps *et al*., 2010; van Velzen *et al*., 2019; Zhao *et al*., 2021).

Subsequently, bacterial symbiosis genes may have been transferred horizontally to other Alphaproteobacteria (Nuti *et al*., 1979; Martínez-Romero, 2009; Yang *et al*., 2020; Rahimlou *et al*., 2021). Conversely, the fewer rhizobia genera in Papilionoideae align with evolutionary events shaping this subfamily’s diversification, potentially leading to a more streamlined nitrogen fixation process as well (Sprent *et al*., 2017; Afkhami *et al*., 2018; Zhao *et al*., 2021).

The use of 16S rRNA metabarcoding represented a valuable tool for studying the microbiome of nodules of legume species where it was not possible to obtain isolates (Hossain *et al*., 2023; Xiang *et al*., 2023). The information generated confirms the large number of bacterial species that coexist in this tissue, thus challenging the old paradigm that nodules are colonized by a single species. However, it did demonstrate the dominance of particular rhizobial species or even the presence of several diazotrophic species. With this analysis, we provide information on the symbionts of many legume species that had not been previously explored or add new data to those with previously existing information.

Finally, in this study, we highlight the importance of studying the legume-rhizobia symbiotic relationships in tropical forest plants, beyond plants of agricultural importance. To our knowledge, few studies of this nature cover such a wide diversity of tropical leguminous forest species. We provide new and unique information on the who with whom and identify multiple rhizobial isolates that may be subject to new species descriptions. We believe that this work contributes significantly to the understanding of the complex relationships between legumes and their rhizobial partners. We strongly encourage further research in this field, encompassing a broader range of species across different legume clades and growth habits worldwide.

## Acknowledgments

Funding was provided by University of Costa Rica (projects B9-204 and C0-524) and by Ministerio de Ciencia, Innovación, Tecnología y Telecomunicaciones de Costa Rica (MICITT)-Consejo Nacional para Investigaciones Científicas y Tecnológicas (CONICIT) (project FI-036B-19). We are deeply grateful for the work that the following plant nurseries are doing to conserve the country’s forest species: Vivero Forestal EDECA, Universidad Nacional; CODEFORSA, San Carlos; Centro de Conservation Santa Ana; Osa Conservation and Asociación Corredor Biológico Talamanca Caribe. We thank botanists Nelson Zamora, Luis G. Acosta, and Stephanie Nuñez for their collaboration in the taxonomic identification of plant species. We also thank Valery Nuñez for her support with the root scanning.

## Author contributions

KRJ designed the research, performed research, data analysis and drafted manuscript; JMH performed research; MBM performed research; BLV performed research; AZO performed research; JMM performed research; LSR performed research; MAB performed research; OVB performed research. All authors commented on and edited the manuscript.

## Data availability

Amplicon sequence data were deposited at the GenBank under accession numbers PP449130-PP449202 (for *mat*K) and PP439182-PP439285 (for the 16S rRNA). Metabarcoding sequence data were deposited at the NCBI Sequence Read Archive under accession number PRJNA1084876 (https://www.ncbi.nlm.nih.gov/sra/PRJNA1084876). Oscar J. Valverde-Barrantes

## Supplementary Material

**Supplementary Fig. 1.**
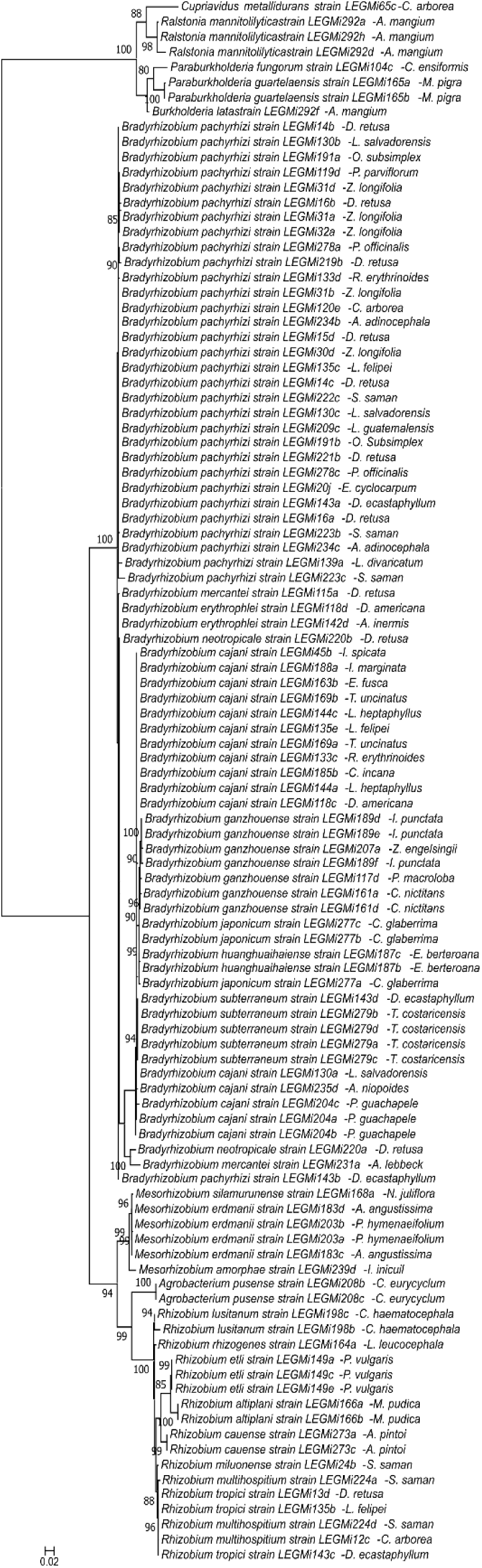
Phylogenetic relationships of the rhizobial strains isolated in this study, based on 16S rRNA sequence dataset was aligned with MAFFT, and the maximum likelihood phylogenetic tree was constructed using IQ-TREE, employing the best-fit models of evolution (TN+F+R3). The resulting trees were visualized and edited in MEGA-X, with. Bootstrap values depicted on the nodes. The name of the best-match species assignation, strain number, and legume host are shown for each strain.

**Supplementary Fig. 2.**
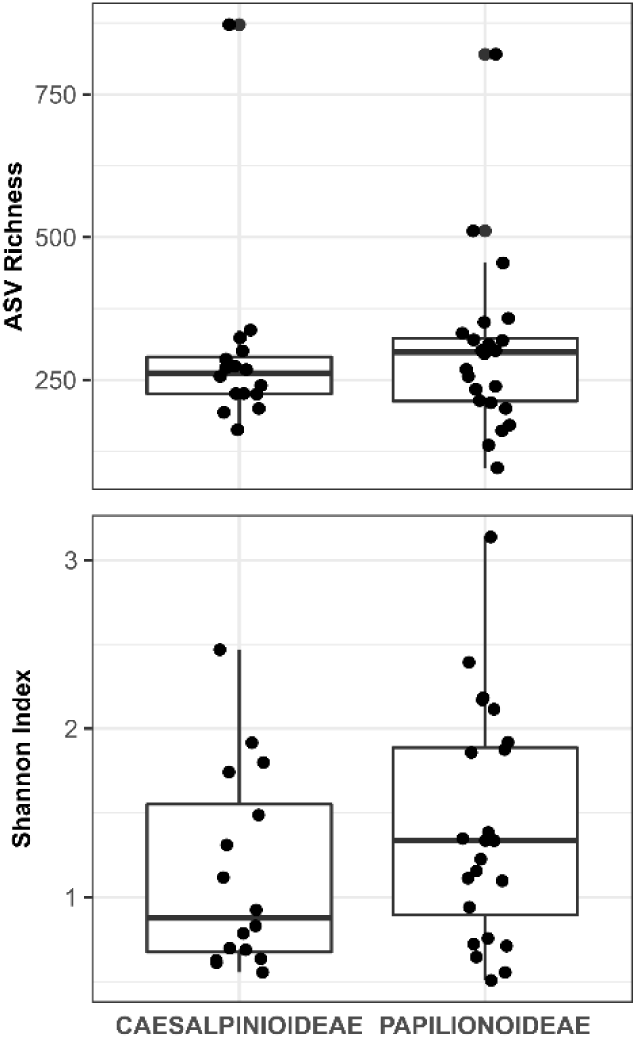
Alpha diversity analyses of the microbial communities in the nodules of sampled legume species in subfamilies Caesalpinioideae and Papilionoideae determined by metabarcoding of the 16S rRNA marker. The upper panel shows the ASV richness values and the lower panel the Shanon index values.

